# Alternate VEGFA isoform expression in fibrosarcoma leads to plasticity in cellular migration and differences in sensitivity to inhibition with anti-VEGFA antibodies

**DOI:** 10.1101/2025.11.09.687017

**Authors:** Yu Chin Lee, Brenda L Aguero Burgos, Nicola H. Green, Frederik Claeyssens, William R. English

**Author notes:** Corresponding Author: William R English, School of Medicine and Population Health, University of Sheffield, S10 2RX.

## Abstract

Metastasis remains a primary contributor to cancer-related mortality. Elevated expression of soluble vascular endothelial growth factor A (VEGFA) isoforms is linked to increased metastatic potential and enhanced cellular plasticity in response to microenvironmental changes. To investigate the connection between VEGFA isoform expression, migration plasticity, and metastasis, we analyzed the migratory behavior of fibrosarcoma cells expressing single VEGFA isoforms within engineered microenvironments mimicking tumor-associated extracellular matrix (ECM) architectures that drive distinct migration modes. Fibrosarcoma cell lines expressing VEGFA120 or VEGFA188 were derived from embryonic (Fs120 and Fs188) or mature skin fibroblasts, ensuring comparable VEGFA isoform expression across cells of different origins. Characterization of collagen fibril diameter and organization in tumors derived from Fs120 or Fs188 cells showed that Fs120 tumors had a higher degree of fibril alignment, suggesting VEGFA120 expression may facilitate ECM remodeling to support metastasis. To simulate these different collagen fibril structures for use in migration studies *in vitro*, we developed 3D platforms representing aligned and randomly organized collagen fibers using electrospinning of polycaprolactone. On 2D surfaces favoring mesenchymal migration and on 3D electrospun scaffolds promoting contact-guided migration, VEGFA120- and VEGFA188-expressing cells exhibited similar migratory capacities, regardless of their origin. However, anti-VEGFA antibodies selectively inhibited the migration of VEGFA120-expressing cells on fibronectin-coated surfaces and on aligned fiber scaffolds, but not non-aligned fibers, revealing isoform-specific sensitivity to VEGFA inhibition. In conclusion, while the expression of single VEGFA isoforms does not directly alter migration plasticity, VEGFA120 expression in mouse fibrosarcomas uniquely promotes ECM reorganization in tumors and cells expressing VEGF120 show increased sensitivity to anti-VEGFA antibodies.

## Introduction

Metastasis is a leading cause of patient mortality, accounting for over 90% of cases [1]. However, its intricate multi-step progression, compounded by factors such as the genetic diversity of cancer cells and microenvironmental influences, enhances adaptability and resistance to therapies, thereby amplifying challenges in prevention and treatment efforts. Vascular endothelial growth factor A (VEGFA), a pivotal regulator of angiogenesis, exhibits three primary isoforms in humans (VEGFA121, VEGFA165, and VEGFA189), attributed to alternative splicing variations [2, 3]. Clinical observations have shown upregulation of VEGFA expression at both mRNA and protein levels across various cancers strongly correlating with tumor growth and metastasis [4]. Elevated VEGFA expression within tumors induces the formation of leaky and enlarged vasculature, facilitating the escape of cancer cells from their primary sites [5]. Besides its role in regulating endothelial cell and vascular biology, VEGFA also influences cancer cell mobility, invasiveness, and chemoresistance [6–9], with disruption of its signaling impairing metastatic traits. Targeting VEGFA with agents like bevacizumab, a function-blocking anti-VEGFA antibody, is a widely used strategy across various cancer types to impede angiogenesis and vascular leakage and increase vascular normalization within tumors, augmenting the action of chemotherapy [10–16]. Yet, the efficacy of these anti-VEGFA therapies in inhibiting metastasis remains poorly understood.

Cell migration in cancer encompasses both collective and individual cellular migration, with individual migration further classified into mesenchymal, amoeboid, and contact-guided modes regulated by the Rho, Rac, and Cdc42 protein families [10, 11]. Mesenchymal migration is initiated by integrin binding, activating Rac and Cdc42 signaling to promote lamellipodia formation, accompanied by the involvement of extracellular proteases [12, 13]. Amoeboid migration is governed by RhoA signaling to ROCK, resulting in actomyosin contraction that facilitates cell squeezing through extracellular matrix (ECM) voids without proteolysis [14]. Contact guidance, also known as 1D migration, refers to the phenomenon wherein cells migrate along narrow linear structures, and it is governed by ROCK signaling [15, 16]. The efficiency of cell movement in contact guidance mode is linked to the arrangement of these linear structures, highlighting the significance of enzymes engaged in ECM modification [17–20]. Successful metastasis relies on cancer cells migrating through diverse microenvironments shaped by the ECM, whose topographical features determine the predominant mode of cell migration. Cancer cells exhibit remarkable plasticity, capable of transitioning between migration modes to adapt to the challenges presented by ECM architecture, thereby sustaining their metastatic potential [13, 21].

The extracellular matrix (ECM) within the tumor microenvironment (TME) plays a pivotal role in either promoting or suppressing tumorigenesis and metastasis by regulating its biochemical components (e.g., growth factors) and biophysical properties (e.g., topography) [22]. Structural changes in the ECM, such as altered cross-linking or porosity, can significantly remodel its architecture within the TME. Key regulatory proteins, including lysyl oxidase (LOX), lysyl oxidase-like proteins (LOXLs), and procollagen-lysine 1,2-oxoglutarate 5-dioxygenases (PLODs), control the formation of fibrillar structures and the stiffness of the ECM [23, 24]. These fibrillar networks function as migratory tracks that facilitate cancer cell movement through the stroma [20]. Concurrently, elevated expression of ECM-degrading proteases enlarges gaps and pores, enhancing cellular penetration [25, 26]. The combined activity of cross-linking enzymes and matrix-degrading proteases thus promotes cancer cell migration and invasiveness, accelerating tumor progression and metastasis across multiple cancer types [27–29]. Given its critical role in disease advancement, ECM architecture within the TME is increasingly evaluated in clinical settings to assess tumor progression and predict treatment responses [30, 31]. This growing recognition has spurred the development of *in vitro* platforms that faithfully recapitulate ECM structure, enabling deeper investigation into the mechanisms driving tumor progression and metastasis, and supporting the innovation of targeted therapeutic strategies.

Our previous studies have shown that differential expression of VEGFA isoforms influences fibrosarcoma metastasis and sensitivity to anti-VEGFA therapy, and the difference in therapeutic response occurs in the primary tumor rather than at the site of metastasis [9]. This led us to hypothesize that differences in VEGFA isoform expression may influence cellular migration within the tumor microenvironment and its sensitivity to anti-VEGFA therapy. In this study, we aimed to address this by utilizing *in vitro* platforms that mimic the ECM architecture observed in the TME of tumors expressing different VEGFA isoforms. We explored the relationship between isoform expression and their plasticity in modes of cell migration using mouse fibrosarcoma cells generated from different cellular origins expressing single VEGFA isoforms. Additionally, we investigated the response of cell migration to anti-VEGFA therapy in these models. Our findings revealed that while the adaptation to various engineered platforms is largely independent of the expression of distinct VEGFA isoforms, only cells expressing VEGFA120 responded to anti-VEGFA therapy. This observation aligns with our previous findings in mouse models of metastasis [9], suggesting the potential of these platforms to investigate mechanisms of metastasis and its inhibition.

## Materials and Methods

### Reagents

The antibody that binds to VEGFA, B20-4.1.1, and its control IgG, BE5, were kindly provided by Genentech. Suramin salt (574625) was purchased from Sigma-Aldrich. Antibodies to SOX2 (ab97959), VEGFR1(ab32152) and GAPDH (ab8245) were purchased from Abcam. NRP1 (3725) antibody was from Cell Signalling.

### Cell culture of fibrosarcoma cell lines

Fibrosarcoma cell lines expressing a single VEGFA isoform (Fs120 and Fs188) were developed from oncogenically transformed and immortalized mouse embryonic fibroblasts isolated from heterozygous breeding pairs [32, 33] as previously described [34]. All VEGFA-expressing cells, as well as transfection control cells, were cultured in Dulbecco’s Modified Eagle’s Medium (DMEM) supplemented with 10% v/v heat-inactivated FBS, 2 mM L-glutamine, 1% penicillin-streptomycin (100 units), 600 µg/mL G418, and 2 µg/mL puromycin. VEGFA knockout mouse fibrosarcoma cells (KO) were kindly provided by Prof. Yihai Cao [35] and were derived from mature skin fibroblasts isolated from Vegfa^+/ +^/Vegfa^-/-^ chimeric mice [36–38]. VEGFA KO cells were cultured in DMEM supplemented with 10% v/v heat-inactivated FBS, 2 mM L-glutamine, and 1% penicillin-streptomycin (100 units).

### Construction of the transposon vector pCLIIP for stable expression of single VEGFA isoforms

The vector containing a pCLIIP backbone harbored a Luciferase2-E2A-mStrawberry-pA fragment (pCLIIP-C-LS), as previously described [9]. The luciferase fragment within the vector was replaced by a synthetic DNA fragment encoding pUC57+ [HRE]_5_-minCMV-VEGFA (Eurofins Scientific) through double digestion with Fse𝙸 and Mlu𝙸 enzymes, thereby yielding the resultant construct, pCLIIP-[HRE]5-minCMV-VEGFA. Initial experiments showed that expression levels of VEGFA120 and VEGFA188 from the mouse *Vegfa* KO fibrosarcoma cells in atmospheric O_2_ using the -[HRE]5-minCMV promoter sequence were significantly lower than expression from the Fs120 or Fs188 cells. We chose to replace the -[HRE]5-minCMV promoter with the CMV promoter sequence. To this end, the enhancer and promoter fragments within the pCLIPP-[HRE]5-minCMV-VEGFA120 and pCLIPP-[HRE]5-minCMV-VEGFA188 vectors were excised using Mlu𝙸 and Not𝙸 enzymes. Subsequently, the CMV promoter from the pcDNA 3.1 vector (Clontech) was amplified using forward 5 ’ - CCGGGGATCTACGCGTGACATTGATTATTGACTAGTTAT -3’ and reverse 5 ’ - CCTACCGGTGCGGCCGCAGCTCTGCTTATATAGACCTC-3’ primers to include a 15-bp sequence recognizing the sticky ends generated post Mlu𝙸 and Not𝙸 digestion. The In-Fusion enzyme (Takara Bio) ligated the digested vector into the CMV promoter, resulting in the pCLIIP-CMV-VEGFA120 and pCLIIP-CMV-VEGFA188 vectors. The integrity and sequence fidelity of all constructed vectors were validated through Sanger sequencing with primers recognizing the CMV promoter and specific VEGFA isoforms.

### Development of adult mouse fibrosarcomas stably expressing single VEGFA isoforms

The vector pCLIIP-CMV-VEGFA120 and pCLIIP-CMV-VEGFA188 were transfected into *mouse Vegfa* KO fibrosarcoma cells seeded in 6-well plates through a mixture with TransIT-X2 and the plasmid PmBP expressing the piggybac transposase. Transfected cells were subsequently identified by their resistance to puromycin. Following transfection, pools of cells were subjected to single colony selection, and colonies exhibiting expression levels of VEGFA comparable to those observed in Fs120 and Fs88 cells were taken forward for further study.

### Characterization of the fibrillar collagen architecture in fibrosarcomas

Paraffin-embedded fibrosarcoma sections from Fs120 or Fs188 tumors were deparaffinized and mounted with PBS, then sealed with clear nail polish. The architecture of fibrillar collagen was imaged using a Zeiss LSM 510 META confocal microscope equipped with 40 x oil lenses. A Chameleon laser, tuned at 940 nm, was focused onto the tumor sections, resulting in second harmonic generation (SHG) signals detected at 470 nm [39]. Images were captured from randomly selected positions using a Chalkey grid in necrotic and viable areas within the same tumor section and analyzed using Orientation J [40].

### Engineering collagen fiber-mimicking *in vitro* microenvironments

The architecture of fibrillar collagen detected in mouse fibrosarcomas was replicated through the electrospinning of polycaprolactone (PCL) dissolved in dichloromethane (DCM). A solution of 15% w/v PCL was loaded into a syringe fitted with a 20-gauge blunt-end needle and pumped at a rate of 1 mL per hour into an electric field with 12.7 kV. Fibers were spun onto a collector covered with foil, positioned 15 cm away from the needle, rotating at 2000 rpm for aligned distribution or 200 rpm for random distribution. The fiber scaffolds were then fastened in CellCrown Inserts (Scaffdex) and washed with sterile PBS before use in subsequent studies.

### Preparation of ECM-coated engineered microenvironments

Cell culture plates were coated with collagen by incubating with rat tail collagen type I (354236 ,Corning) at a concentration of 20 µg/mL in PBS at room temperature for 2 hours. The fibronectin layer on the culture plate was made by culturing a solution of fibronectin (FC010, Merck) at a concentration of 5 µg/mL in PBS at room temperature for 2 hours. For coating fiber scaffolds with fibronectin, the scaffolds were immersed in a solution of fibronectin at a concentration of 10 µg/mL in PBS and left overnight at room temperature on a shaker. Following the coating process, the plates or fibers were washed twice with sterile PBS and rinsed with a culture medium before cell seeding.

### Immunofluorescence staining of cells on fiber scaffolds

Cells were seeded at a density of 5×10^4^ cells in 500 µL of the medium onto pre-fibronectin-coated fiber scaffolds and cultured overnight. Cells were fixed with 4% w/v PFA in PBS and permeabilized with 0.1% v/v Triton X-100 in PBS supplemented with 1% w/v BSA, subsequently blocked with 1% w/v BSA in PBS. Cells were stained with a primary antibody against fibronectin (Abcam, ab2413) at a dilution of 1:200 and incubated at room temperature for 1 hour in the dark. Following primary antibody incubation, cells were incubated with a secondary antibody conjugated with Alexa Fluor 488 (Invitrogen) at a dilution of 1:500 for an hour at room temperature in the dark. After antibody staining, cells were further stained with Phalloidin (1:400) at room temperature for 30 minutes in the dark. DAPI staining was performed overnight in the dark.

### Time-lapse single-cell migration assay

Cells pre-seeded at a density of 1×10^4^ cells on 2D ECM-coated surfaces or at a density of 4×10^4^ cells on fibronectin-coated fiber scaffolds were subjected to serum starvation in 500 µL of DMEM supplemented with 2 mM L-glutamine, 1% penicillin-streptomycin (100 units), and either 40 µg/mL of anti-VEGFA antibody B20-4.1.1 or its control IgG for 2 hours. A chemotactic gradient of the growth factor was established using 10% v/v horse serum (H01146, Sigma-Aldrich) in low-gelling temperature agarose dissolved in PBS, placed on one side of the plates 30 minutes before the initiation of time-lapse imaging. Throughout the imaging process, cells were maintained at 37 ℃ in an atmosphere supplemented with 5% CO_2_. Images of cells at multiple positions were captured every 10 minutes for 3 hours using a Zeiss CellDiscoverer 7 microscope equipped with a 10 x lens. The locations of 30 randomly selected cells at various time points were tracked using the Cell Tracking tool in ImageJ. Subsequently, the migration of each cell line was quantified using the Chemotaxis Tool provided by Ibidi.

### Statistical analysis

The data were plotted and analyzed using GraphPad Prism 9 (GraphPad Prism Software Inc.). Parametric grouped data were analyzed using the ANOVA test followed by multiple comparisons with Fisher’s LSD test. In contrast, non-parametric data were analyzed using the Kruskal-Wallis test, followed by multiple comparisons with the Uncorrected Dunn’s test. Significant differences between selected comparison groups were indicated as described in the figure legends if they were present using *p < 0.05, *p < 0.05, ** p < 0.01, and *** p < 0.001.

## Results

### Fibrosarcoma cells derived from different murine cellular origins were engineered to express single VEGFA isoforms at similar levels

Fibrosarcoma cells expressing specific single murine VEGFA isoforms were sourced from two distinct origins: embryonic fibroblasts isolated from heterozygous breeding pairs and mature VEGFA knockout skin fibroblasts from chimeric mice. The variations between cellular sources generated cells with unique developmental backgrounds. Figures 1 (A) and (B) illustrate the developmental processes leading to the generation of VEGFA120- and VEGFA188-expressing fibrosarcoma cells derived from both embryonic fibroblasts (Fs120 and Fs188) [34] and mature skin fibroblasts (VEGFA120 and VEGFA188), along with their identification procedures. While fibrosarcoma cells derived from embryonic fibroblasts express VEGFA isoforms from the endogenous mouse *Vegfa* promoter, those derived from mature skin fibroblasts have stable transfection for isoform expression from a plasmid under the control of an artificial promoter. Based on the pCLIIP backbone, the transfection vector was constructed to carry genes encoding specific VEGFA isoforms in conjunction with a CMV promoter (Figure S1) [9]. The production of total VEGFA protein, as measured by ELISA in the conditioned medium collected from the cells, showed significantly higher levels of secretion from Fs188 cells than from Fs120 cells. Furthermore, the protein expression of VEGFA from VEGFA knockout fibrosarcoma cells was comparable to levels detected from Fs120 or Fs188 cells. In contrast, no VEGFA was detected in the medium collected from VEGFA knockout fibrosarcoma cells transfected with pCLIIP alone (Figure 1C).

**Figure 1.**
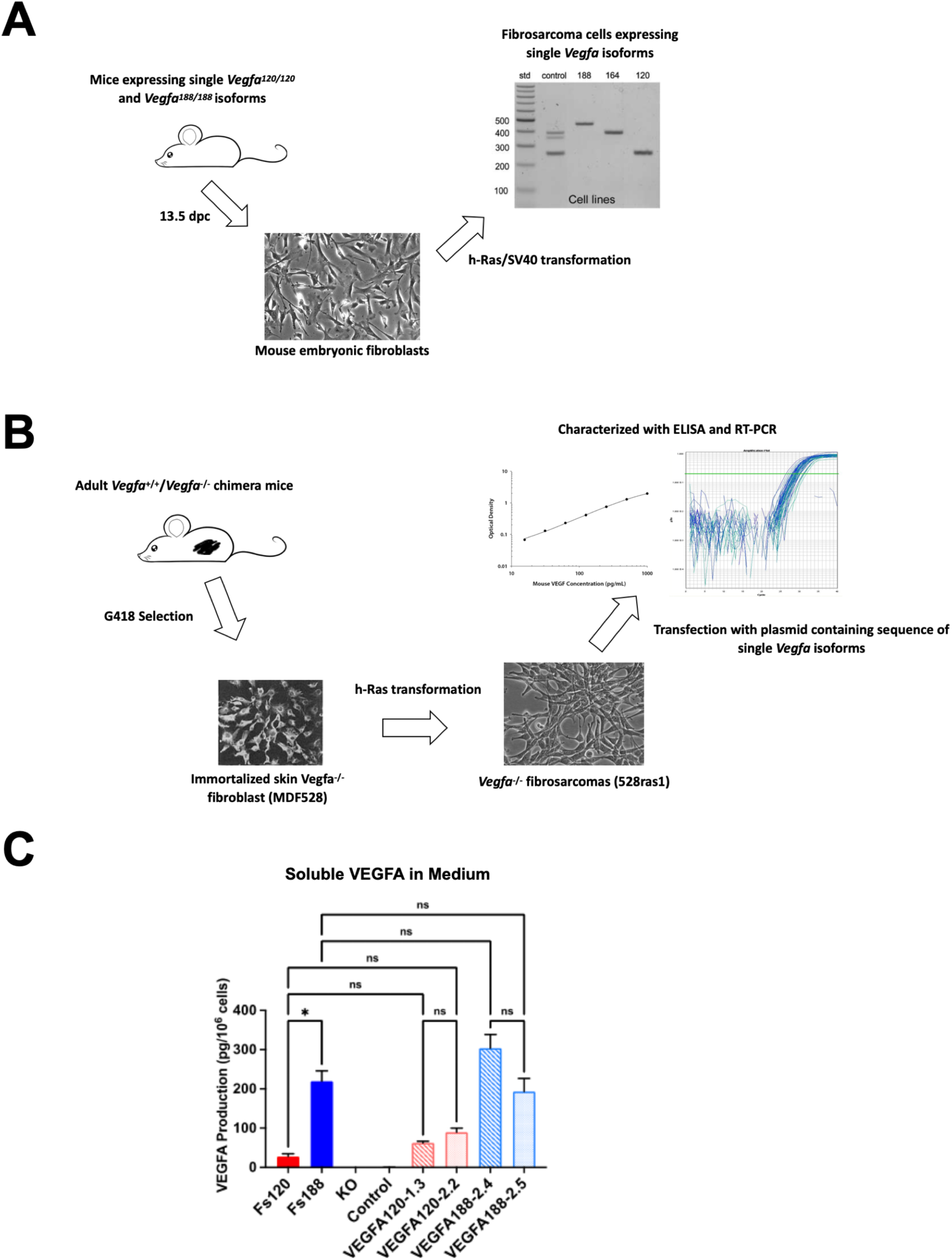
Steps for developing fibrosarcoma cells expressing a single VEGFA isoform from **(A)** embryonic fibroblasts isolated from heterozygous breeding pairs and **(B)** mature skin total VEGFA knockout fibroblasts. **(C)** ELISA measurements of soluble VEGFA protein expression in cultured media collected from different fibrosarcoma cell lines. Each column represents the mean of three independent replicates, with error bars denoting ± SEM. Statistical significance (*p < 0.05) and non-significance (ns) were determined using the non-parametric Kruskal-Wallis test, followed by multiple comparisons using an Uncorrected Dunn’s test.

### The migration of fibrosarcoma cells on 2D ECM-coated surfaces is independent of cellular origins but shows isoform-selective sensitivity to anti-VEGFA treatment

Investigations into cell migration predominantly start with those on 2D surfaces due to their easily reproducible characteristics and adaptability to various experimental conditions. Figure 2 (A) illustrates the setup of the 2D-engineered microenvironment used in our studies, featuring a thin layer of ECM proteins supplemented with a chemotactic gradient created by horse serum. Unlike the commonly employed wound-healing (scratch) assays, which primarily assess collective cell migration and lack chemoattractant cues, our *in vitro* platform enables the characterization of single-cell migration initiated by chemoattractant. Despite their diverse cellular origins, fibrosarcoma cells expressing single VEGFA isoforms showed no significant differences in migration speed on surfaces coated with collagen-I (Figure 2B). However, significant variations in migration speed were observed among cells originating from different cellular sources. Specifically, cells derived from mature skin fibroblasts exhibited faster migration on fibronectin-coated surfaces compared to embryonic fibroblasts (Figure 2C). Moreover, we investigated the dependence of cell migration on VEGFA by using B20-4.1.1, an engineered anti-mouse VEGFA monoclonal antibody designed to mimic bevacizumab for pre-clinical murine studies [41]. The inhibition of migration by B20-4.1.1 was only observed in VEGFA120-expressing fibrosarcoma cells on fibronectin-coated surfaces and was independent of cellular origin (Figure 2C).

**Figure 2.**
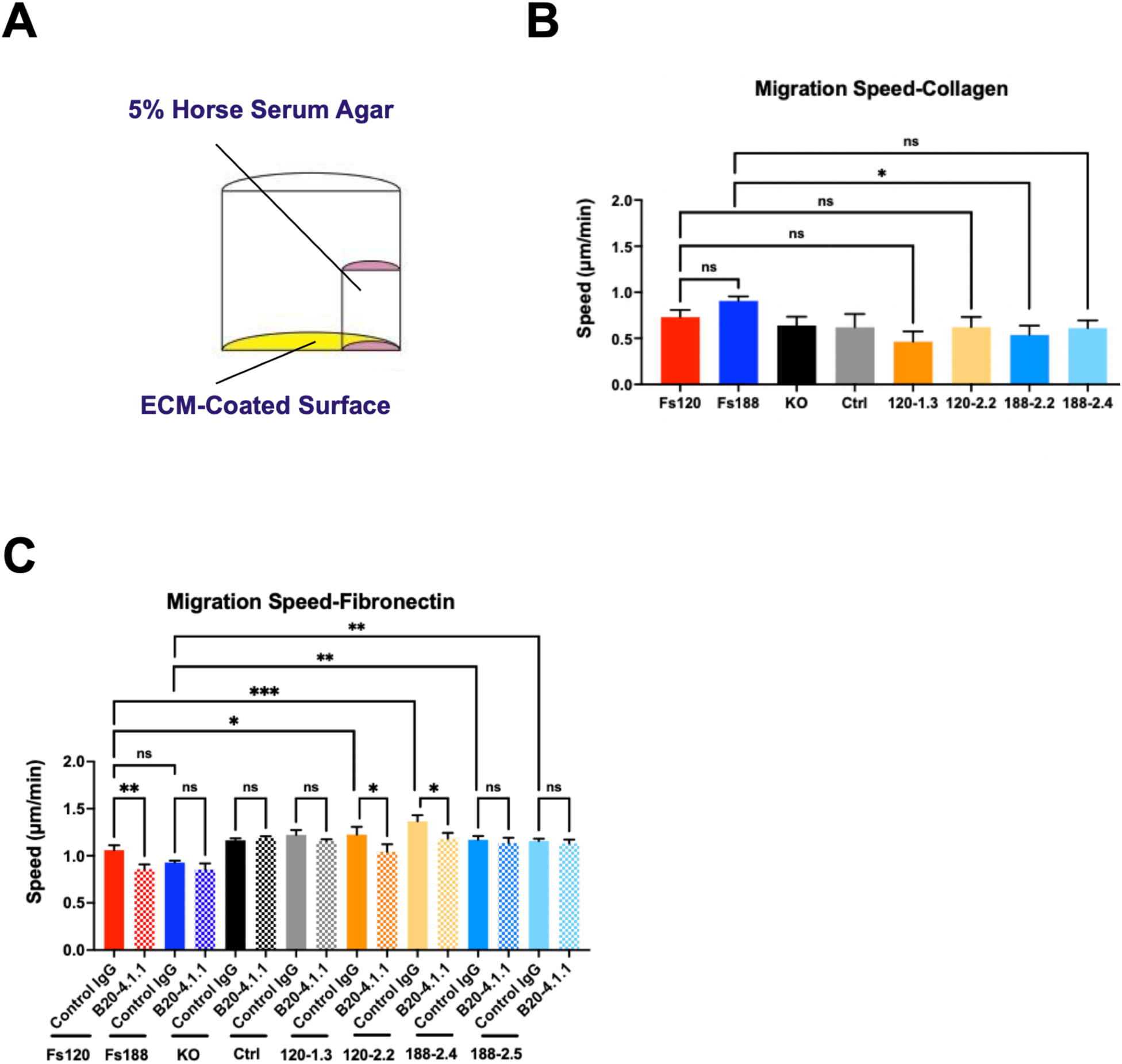
**(A)** The design of an *in vitro* platform with a chemotactic gradient, facilitated by 5% horse serum in agar, on 2D surfaces coated with ECM protein. **(B)** Quantification of migration speed across fibrosarcoma cell lines on collagen-coated surfaces from time-lapse imaging. **(C)** Quantification of migration speed across fibrosarcoma cell lines on fibronectin-coated surfaces from time-lapse imaging in the presence of 40 µg/mL B20-4.1.1 or its control IgG. Each column represents the mean of three independent replicates, with error bars denoting ±SEM. Statistical significance (*p < 0.05, ** p < 0.01, *** p < 0.001) and non-significance (ns) were determined using **(B)** the non-parametric Kruskal-Wallis test with multiple comparisons by Uncorrected Dunn’s test or **(C)** the ANOVA test with multiple comparisons by Fisher’s LSD test.

### Electrospinning of PCL fiber scaffold mimics the architecture of collagen fibers derived from mouse fibrosarcomas expressing single VEGFA isoforms

Unlike the simplistic model of 2D ECM-coated surfaces, the ECM architecture in the TME is considerably more complex. Developing novel *in vitro*-engineered microenvironments that recapitulate the characteristics of ECM architecture in the TME could advance our understanding of the molecular mechanisms that regulate metastasis. H&E staining revealed the histological features of viable and necrotic regions in tumor sections derived from either Fs120 or Fs188 cells (Figure 3A). These distinct features served as references for guiding the acquisition of SHG images. Figure 3B shows examples of SHG signals generated by collagen fibers in both viable and necrotic regions. Using the hue-saturation-brightness (HSB) color-coding scheme, which assigns distinct colors to oriented structures based on their angles, collagen fibers with different orientations were visualized in their corresponding colors. In the viable regions of Fs120 tumors, collagen fibers were predominantly coded in uniform colors, indicating a consistent orientation relative to the reference horizon. In contrast, fibers in the viable regions of fs188 tumors, as well as in the necrotic areas of both tumor types, exhibited random coloration, reflecting a distribution across multiple angles (Figure 3C). These findings suggest that collagen fibers were more aligned in the viable regions of Fs120 tumors, whereas fibers in the viable regions of Fs188 tumors and the necrotic regions of both Fs120 and Fs188 tumors displayed a random orientation. Moreover, in the viable regions, collagen fibers in Fs188 tumors demonstrated a significant increase in diameter compared to those in Fs120 tumors. However, the number of branching points in Fs188 tumors was notably reduced (Figure 4A). Despite the significant differences in fiber diameter between Fs120 and Fs188 tumors, the diameter consistently ranged between 2 to 3 µm (Figure 4A). By optimising the syringe pump speed for PCL fiber desposition, we successfully created electrospinning fiber scaffolds with different fiber orientations that replicated the fiber diameter of between 2 to 3 µm seen *in vivo* (Figure 4B) and allowed cells to attach after coating with fibronectin (Supplementary Figure S2).

**Figure 3.**
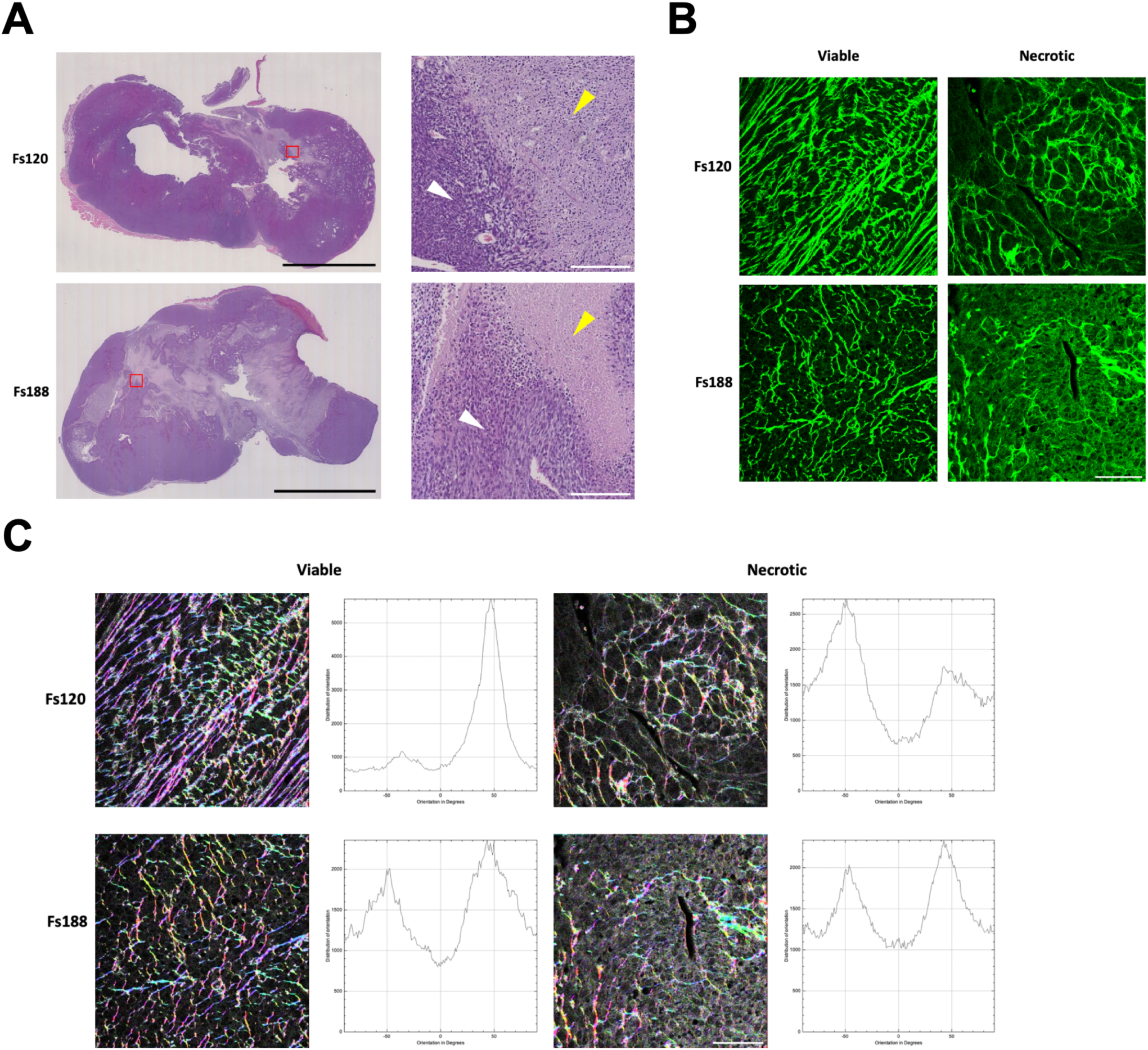
**(A)** Hematoxylin and eosin staining of paraffin-embedded tumor sections from fs120 and fs188 tumors distinguished viable (white arrow) or necrotic (yellow arrow) regions. Black scale bar, 1 cm. White scale bar, 200 µm. **(B)** Representative second-harmonic images of fibrillar collagen architecture (depicted in green) in tumor sections between viable and necrotic regions were acquired under 40 x oil lenses using a Zeiss LSM 510 META confocal microscope with a Chameleon laser tuned at 940 nm excitation and signal collected at 470 nm. Scale bar, 50 µm. **(C)** Representative HSG color-coded maps illustrate the orientation of collagen fibers in tumor sections. According to the angles to the reference horizon, fibers were coded in different colors. Fibers with similar orientations were color-coded the same, and their orientation distribution was plotted based on their tilted angle. Scale bar, 50 µm.

**Figure 4.**
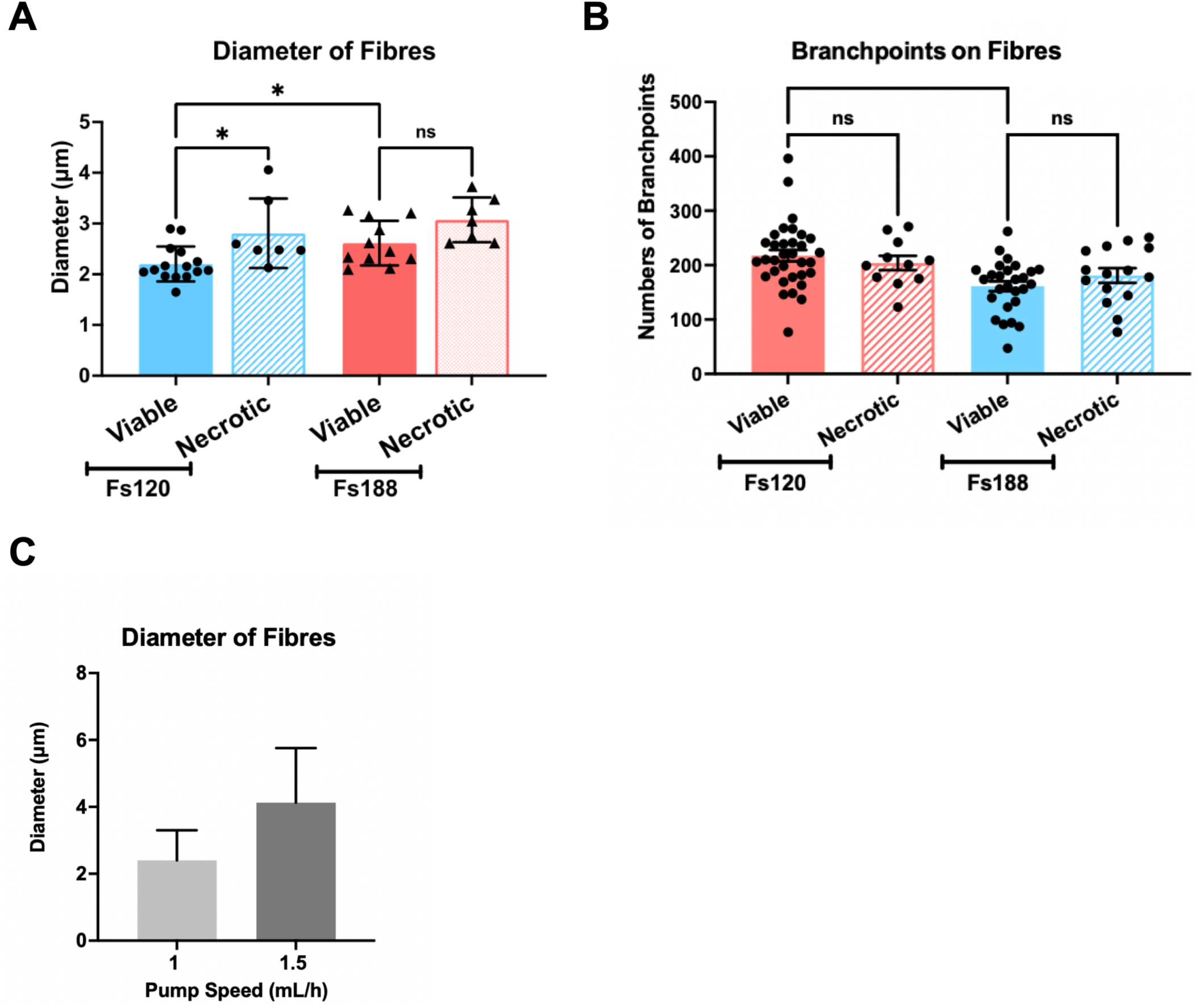
**(A)** Quantification of collagen fibers’ diameter and **(B)** branching points in tumor sections between viable and necrotic regions. Each column represents the mean, with error bars denoting ±SEM (Fs120/viable, n = 15; Fs120/necrotic, n = 7; Fs188/viable, n = 11; Fs188/necrotic, n = 7). Statistical significance (*p < 0.05, ** p < 0.01, *** p < 0.001) and non-significance (ns) were determined using the non-parametric Kruskal-Wallis test with multiple comparisons by Uncorrected Dunn’s test. **(C)** Quantification of the diameter of electrospun fibers produced under different pump speeds. Each column represents the mean of 300 measurements from 3 images, with error bars denoting ±SEM.

### The inhibition of migration of fibrosarcoma cells on electrospun fiber scaffolds is dependent on the difference in expression of VEGFA isoforms

Figure 5 (A) illustrates the configuration of the engineered 3D *in vitro* platforms, where fibronectin-coated fiber scaffolds were anchored at the bottom of the well plate using cell crown inserts. The differences in migration speed seen between fibrosarcoma cells expressing distinct VEGFA isoforms and originating from different cellular sources showed no significant differences between aligned or randomly orientated fibers (Figure 5B, C). However, the inhibitory effects of B20-4.1.1 on migration in VEGFA120-expressing fibrosarcoma cells were only observed on aligned fiber scaffolds, not randomly orientated scaffolds (Figure 5B). Similar to the observation on 2D surfaces, VEGFA188-expressing cells remained resistant to B20-4.1.1 inhibition of migration on both types of fiber scaffolds. The migration behavior observed when cells move along fibers is commonly referred to as “contact guidance,” wherein the orientation of fibers strongly influences the direction of migration [42, 43]. Trajectory plots, showing the migration paths of fibrosarcoma cells, showed migration direction aligned with the orientation of the fibers and was independent of VEGFA isoform expression and tissue origin of the cells (Figure 5D). Furthermore, cells exhibited randomly distributed migration paths across multiple angles when migrating on randomly orientated fibers (Figure 5E). Although the chemotactic gradient was used, fibrosarcoma cells did not migrate toward the chemoattractant, indicating that chemotaxis is not the major determinant of direction while migrating on fibers in our model system (Figure 5D, E).

**Figure 5.**
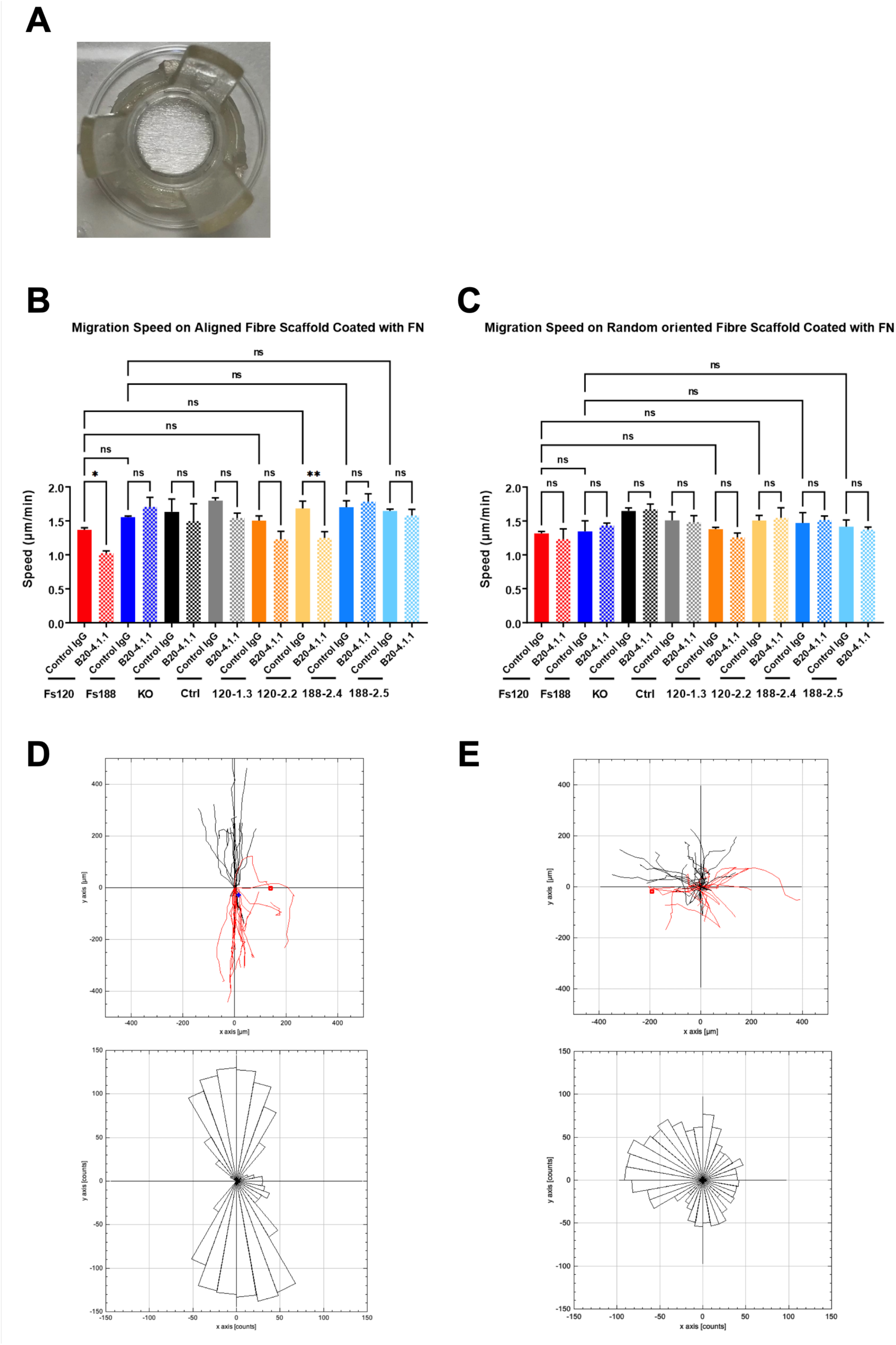
**(A)** Engineered 3D *in vitro* platforms were created to mimic the fibrillar collagen architecture in the TME by fastening the electrospinning fiber scaffolds in the CellCrown Insert. **(B-C)** Quantification of migration speed across fibrosarcoma cell lines was conducted on fibronectin-coated **(B)** aligned and **(C)** random-orientated fiber scaffolds using time-lapse imaging in the presence of 40 µg/mL B20-4.1.1 or its control IgG. Each column represents the mean of three independent replicates, with error bars denoting ±SEM. Statistical significance (*p < 0.05 and ** p < 0.01) and non-significance (ns) were determined using the non-parametric Kruskal-Wallis test with multiple comparisons by Uncorrected Dunn’s test. **(D-E)** Representative trajectory and rose plots depicting the migration paths and directionality of fibrosarcoma cells on (D) aligned and (E) random-oriented fiber scaffolds.

## Discussion

In this study, we demonstrate that the differential expression of VEGFA isoforms plays a pivotal role in regulating fibrosarcoma cell migration and modulating ECM architecture, providing valuable insights into the mechanisms driving tumor metastasis. By utilizing an advanced model system, we were able to investigate the migratory behavior of mouse fibrosarcoma cells expressing single VEGFA isoforms on distinct *in vitro* platforms designed to mimic various ECM architectures. This approach allowed us to more effectively unravel the relationship between the plasticity of cell migration modes and the endogenous expression of specific VEGFA isoforms. Although clinical strategies aim to target VEGFA to control angiogenesis [44], their effectiveness in inhibiting metastasis remains limited. Our previous studies have shown reduced metastasis in mice with VEGFA120-expressing tumors treated with anti-VEGFA therapy, with this isoform-selective effect identified as originating in the primary tumor rather than at the site of metastasis [9]. In line with these findings, our data show that fibrosarcoma cells expressing VEGFA120 exhibit reduced migratory capacity when treated with anti-VEGFA therapy, in contrast to cells expressing other isoforms. This suggests that VEGFA isoform-specific signaling may be a key factor promoting metastasis and influencing therapeutic response. Moreover, we observed that VEGFA120 expression influences ECM remodeling, as collagen fibrils have more organized, aligned structures seen in tumors derived from VEGFA120-expressing fibrosarcomas compared to those expressing VEGFA188. These results suggest that elevated VEGFA120 expression could serve as a potential biomarker for assessing metastatic risk and optimizing anti-VEGFA treatment strategies in fibrosarcoma.

Fibrosarcoma cells expressing VEGFA120 (Fs120) also exhibit greater plasticity compared to those expressing VEGF188 (Fs188), as evidenced by their distinct morphologies on different ECM-coated surfaces and their ability to migrate using either mesenchymal or amoeboid movement, depending if they are on 2D surfaces or navigating through non-adherent confined gaps (Figure S3) [9, 45]. These findings suggest that the capacity of fibrosarcoma cells to adapt to their migration to various microenvironments may be influenced by differential VEGFA isoform expression. However, VEGFA isoforms are known to have distinct functions during embryonic development [46–48], raising the possibility that fibrosarcoma cells derived from fibroblasts isolated from single VEGFA isoform-expressing embryos may have undergone different levels of selection and differentiation before oncogenic transformation. As a result, it remains unclear whether the observed plasticity is directly linked to VEGFA isoform expression or influenced by differences in cellular differentiation. Given that VEGFA can induce the expression of stem cell markers such as SOX2, which promotes EMT, metastasis, and self-renewal, we generated new cell lines from mature skin fibroblasts to distinguish the effects of VEGFA isoforms from those of cellular differentiation status [49, 50]. These new lines, which lack SOX2 expression and share identical developmental backgrounds, displayed similar phenotypes on different ECM surfaces as the original fibrosarcoma cells (Fs120 and Fs188) (Figure S4). This suggests that isoform-specific signaling primarily regulates the observed differences in migratory capacity and adaptability.

Plasticity in cell migration modes is essential for successful metastasis because different ECM architectures present unique physical challenges [51–53]. Dense, rigid ECMs often require mesenchymal migration, which depends on integrin-mediated adhesion and proteolytic degradation of the matrix. In contrast, softer or less organized ECMs support faster amoeboid movement, are less reliant on matrix remodeling [13, 16, 54]. This ability to switch between migration modes allows cells to adapt to the diverse ECM environments encountered during metastasis. Our study found that mesenchymal migration capacity on 2D ECM-coated surfaces was not influenced by VEGFA isoform expression (Figure 2B and 2C). This contrasts with previous findings where VEGFA188-expressing fibrosarcoma cells were reported to migrate faster than VEGFA120-expressing cells on 2D surfaces [9, 45]. The discrepancy is likely due to the absence of a chemotaxis gradient in previous experiments, emphasizing the importance of chemical cues in initiating migration. Provenzano et al. demonstrated that the reorganization of the fiber architecture controls contact-guided migration, a process driven by actomyosin contractions mediated through Rho/ROCK signaling [42]. Although aligned fibers enhance migration efficiency, we observed no significant difference in contact-guided movement between VEGFA120- and VEGFA188-expressing cells on aligned or randomized fiber scaffolds (Figure 4B and 4C) [17, 43]. This suggests that the ability to migrate on pre-aligned fibers is independent of VEGFA isoform expression. However, our randomized scaffolds may have needed to be sufficiently disordered to challenge the cells’ capacity for fiber reorganization (Figure 3B and 3C), leaving uncertainty about whether VEGFA isoforms modulate fiber remodeling [43]. Additionally, Fs120 cells demonstrated superior amoeboid migration when constrained between two non-adherent surfaces compared to Fs188 cells (Figure S3), which aligns with Kanthou et al.’s findings that Fs120 cell migration relies on actomyosin contractility [45]. These results suggest that VEGFA120-expressing cells exhibit greater plasticity in migration modes, though the most significant differences between VEGFA isoform-expressing cells were primarily observed in amoeboid movement. Despite showing similar contact-guided migration capacity on engineered fiber scaffolds, VEGFA188-expressing cells formed tumors with more disorganized ECM *in vivo*, reducing migration efficiency. These findings are consistent with reports that metastasis increased from tumors derived from VEGFA120-expressing cells in mice, demonstrating higher metastatic potential and further underscoring VEGFA’s role in regulating plasticity in cell migration modes and its links to metastatic potential [9].

Distinct dependencies on integrin adhesion and actomyosin contractility characterize amoeboid and mesenchymal migration [16, 55]. In mesenchymal migration, Rho/ROCK-driven contractility is crucial, facilitating the formation of focal adhesions and stress fibers that generate the traction necessary for forward movement [14, 56]. Conversely, amoeboid cells display weaker integrin-ECM adhesion and depend predominantly on cortical myosin II contractions to navigate through 3D environments [57]. Notably, inhibiting Rho/ROCK or actomyosin contractility impairs amoeboid migration, while having minimal impact on mesenchymal movement, indicating that contractility is vital for amoeboid motility but less critical for mesenchymal migration [21, 58]. Interestingly, Kanthou et al. revealed that only VEGFA120-expressing cells on plastic acquired a mesenchymal phenotype and enhanced migration capacity in response to Rho/ ROCK inhibition [45]. This finding suggests that the reliance on Rho/ROCK signaling in mesenchymal migration differs between VEGFA isoform-expressing cells. Furthermore, VEGFA isoforms are known to regulate migration through activating Rho/ROCK signaling [59]. This raises the interesting possibility that the differences in response to anti-VEGFA therapy of cell migration between VEGFA isoform-expressing cells is linked to Rho/ROCK signaling. In our study, anti-VEGFA therapy selectively inhibited the migration of VEGFA120-expressing cells on fibronectin-coated surfaces (Figure 2C). This selective inhibition likely stems from the strong adhesion of these cells to fibronectin; when Rho/ ROCK-mediated detachment at the cell’s rear is lost, movement is severely impeded [60]. In contrast, VEGFA120-expressing cells on collagen, which exhibit weaker adhesion, showed no response to anti-VEGFA therapy, possibly because their migration does not depend as heavily on Rho/ROCK signaling [61]. Moreover, our experiments using non-adherent chambers, which force cells to adopt amoeboid movement, demonstrated that blocking VEGFA120 with anti-VEGFA antibodies impaired the cells’ ability to navigate confined gaps, consistent with previous observations (Figure S3). In addition to generating the force required for migration, Rho/ROCK-regulated actomyosin contraction also plays a crucial role in organizing ECM fiber structures, which is essential for contact-guided migration [42], which may also provide an explanation for the differences in collagen fiber alignment in the Fs120 tumors. Overall, one explanation for the differential response to anti-VEGFA therapy between VEGFA120- and VEGFA188-expressing fibrosarcoma cells could be linked to differences in their dependence on Rho/ROCK signaling. Further investigations are needed to identify the mechanism by which the different VEGFA isoforms modulate migration modes and the relative dependence on VEGF receptors and integrins that may connect to Rho/ROCK signaling [62–65].

VEGFA isoforms orchestrate extracellular matrix (ECM) reprogramming within the tumor microenvironment (TME) through distinct interactions with VEGF receptors and co-receptors [66]. In colorectal cancer liver metastases, inhibition of VEGF receptor signaling leads to the accumulation of hyaluronic acid and sulfated glycosaminoglycans, increasing tissue stiffness [67]. In contrast, elevated VEGFA levels in breast cancer remodel the ECM by reducing collagen fibers, fibronectin, and hyaluronan, while upregulating MMP1, uPAR, and LOX and downregulating MMP2 and ADAMTS1, illustrating how VEGFA-driven signaling can shape ECM composition and architecture [68]. Our fibrosarcoma models further demonstrate isoform-specific reprogramming: VEGFA120 enhances laminin deposition and induces aligned, linear collagen fibers, whereas VEGFA188 promotes a collagen-rich but disorganized matrix (Figure 3) [9]. This difference in collagen architecture may result from altered expression of procollagen-processing enzymes and collagen-modifying proteins, influenced by isoform-specific downstream signaling [69, 70]. While VEGFR1 and NRP1 are constitutively expressed in these models, VEGFR2 is absent at baseline but becomes inducible by VEGFA in cells expressing single isoforms, indicating that VEGFA-driven ECM remodeling predominantly occurs through the VEGFR1/NRP1 axis (Figure S5) [45]. Crucially, VEGFR1 auto-phosphorylation at Tyr1213 activates Erk1/2 signaling to promote tumor invasiveness, and inhibition of this phosphorylation suppresses the fibrotic phenotype [71, 72]. These data suggest that VEGFA isoforms can fine-tune ECM composition and architecture by modulating the expression of key matrix-regulating enzymes via VEGFR1/NRP1 signaling. Furthermore, the resulting changes in ECM stiffness and fiber alignment are sensed by collagen-binding integrins—particularly α1β1 and α2β1—which co-localize with VEGFR1/NRP1 complexes and amplify downstream FAK and PI3K/Akt signaling [73–76]. This signaling convergence on Erk1/2 further enhances invasive behavior within aligned, stiffened matrices. Collectively, these findings underscore VEGFR1-mediated ECM remodeling—and its functional crosstalk with integrin pathways—as a central mechanism in tumor progression and a promising therapeutic target [77, 78].

Overall, we conclude that fibrosarcoma cells expressing VEGFA120 have more plasticity than those expressing VEGFA188 in utilizing various migration modes to adapt to engineered microenvironments that mimic the diverse ECM architectures encountered during metastasis. However, while anti-VEGFA therapy selectively inhibits the migration capacity of VEGFA120-expressing cells, this does not affect all migration modes. Further work is needed to elucidate the mechanisms that regulate this, with Rho/ROCK being of particular interest.

In summary, our study provides new insights into the role of distinct VEGFA isoforms in regulating cell migration plasticity. We demonstrate that VEGFA120 expression is associated with enhanced adaptability to diverse artificial environments, and ECM architectures and their modification. In contrast, VEGFA188 expression appears to restrict cells to mesenchymal motility. These findings highlight the functional diversity of VEGFA isoforms in driving different modes of cell migration, including mesenchymal, amoeboid, and contact guidance, all of which are critical for tumor invasion and metastasis. Given the role of individual VEGFA isoforms in regulating these migration modes, their distinct responses to anti-VEGFA therapy, and their capability to modify ECM architecture, our results support previously observed isoform-specific therapeutic effects in mice. This work may aid in identifying novel biomarkers and downstream signaling pathways, offering specific therapeutic targets to inhibit key processes driving metastasis. Furthermore, these newly engineered *in vitro* platforms, which are replicable and versatile, provide valuable opportunities to dissect the molecular mechanisms underlying various migration modes and offer robust tools for drug testing, aligned with the 3Rs principle. Moving forward, although anti-VEGFA therapy has primarily targeted abnormal angiogenesis, a deeper exploration of VEGFA isoform-specific effects on migration plasticity and ECM modification could extend its application to more effectively target metastasis.

## Acknowledgements

YCL was funded by a Taiwanese Government Scholarship to Study Abroad. BLAB was supported by the Weston Park Hospital Cancer Charity, UK and a scholarship from CONACyT, Mexico. C, N We would like to thank Genetech^TM^ for the provision of the B20-4.1.1. and control antibodies and Professor Yi Hai Cao, Karolinska Institutet, Sweden for the VEGFA KO cells used in this study. We would also like to thank Claudia Madrigal Esquivel and Nada Al Shaaili for their help with PCR and cloning. Maggie Smith for assistance with histology, Caitlin E Jackson, and Mina Aleemardani for their assistance with plasma coating and electrospinning.

Author Contributions:

