## Supplemental Materials and Methods for "Alternate VEGFA isoform expression in fibrosarcoma leads to plasticity in cellular migration and differences in sensitivity to inhibition with anti-VEGFA antibodies"

### Supplementary Materials and Methods

#### Immunoblotting

Cancer cell proteins were extracted using RIPA lysis buffer supplemented with 5 mM EDTA, protease inhibitors, and phosphatase inhibitors. Protein lysates (30 μg per sample) were mixed with 10 μL of Laemmli buffer containing 10% (v/v) DTT and heated at 95°C for 5 minutes. The proteins were then separated on 10% SDS-PAGE gels and transferred onto a nitrocellulose membrane. The membrane was incubated with primary antibodies (1:1000), followed by secondary antibodies (1:10,000). Protein detection was performed using enhanced chemiluminescence (ECL) reagents.

#### Preparation of confined non-adherent chamber for migration studies

Square glass coverslips (22 mm × 22 mm) were hydroxylated in Piranha solution for 90 minutes at room temperature, followed by two rinses with distilled water for 5 minutes each. The coverslips were then silanized with 3-aminopropyl triethoxysilane under vacuum for 30 minutes and annealed in an oven at 110°C for 30 minutes. To enhance polymer binding, the treated surfaces were crosslinked with 0.5% (v/v) glutaraldehyde in distilled water for 30 minutes. Next, the coverslips were coated with a mixture of 5.6% (w/v) polyethylene glycol (PEG) in distilled water and 2 M hydrochloric acid for 4–5 minutes at room temperature. The coating was evenly distributed using a spin coater set to 7000 rpm with an acceleration of 550 rps/s for 40 seconds. The resulting non-adherent glass coverslips were stored at 4°C until use.

Before the experiment, non-adherent coverslips were rinsed twice with distilled PBS and once with serum-free medium. The bottom coverslip was secured onto a slide with the PEG-coated surface facing up. Cells were seeded at a density of 5 × $10^3$ cells in 100 μl of serum-free medium onto this surface, then covered with another PEG-coated coverslip facing down, separated by a 40 μm-thick layer of 3M Scotch tape. To establish a chemotactic gradient, 10% (v/v) horse serum in low-gelling temperature agarose dissolved in PBS was added between two sets of confined non-adherent coverslip sandwiches 30 minutes before the start of time-lapse imaging (Figure S3A). The confined non-adherent chamber system was then ready for the single-cell migration assay.
