## Supplemental Figure 1 for "Alternate VEGFA isoform expression in fibrosarcoma leads to plasticity in cellular migration and differences in sensitivity to inhibition with anti-VEGFA antibodies"

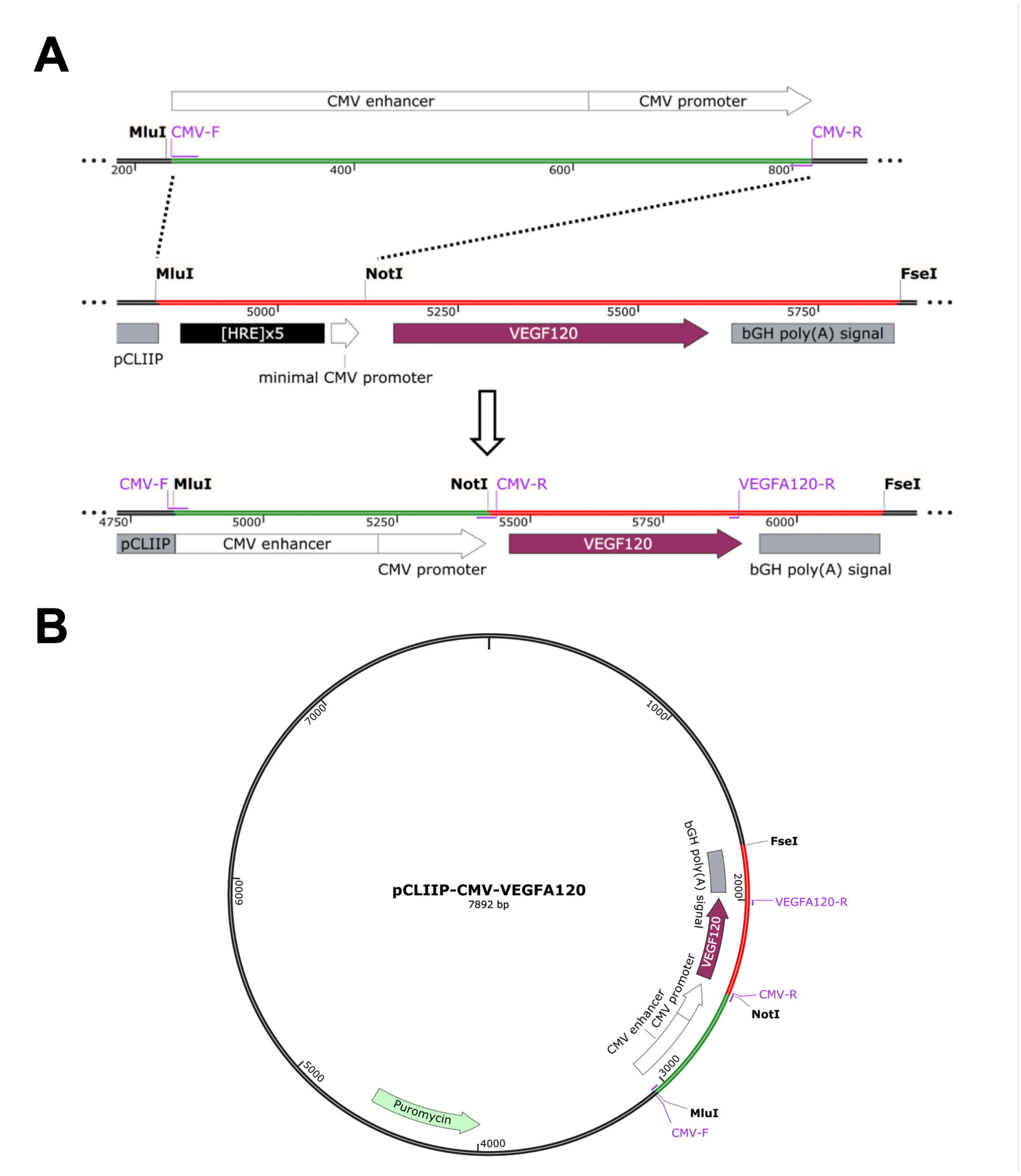


**Figure S1. (A)** Representative image illustrating the replacement of [HRE]$_5$-minCMV with CMV using designed primers that recognize digestion sites of MluI and NotI **(B)** A representative vector containing sequences encoding the CMV promoter, a single VEGFA isoform, and puromycin transfects total VEGFA knockout fibrosarcoma cells, enabling the continuous expression of VEGFA isoforms.
