## Supplemental Figure 2 for "Alternate VEGFA isoform expression in fibrosarcoma leads to plasticity in cellular migration and differences in sensitivity to inhibition with anti-VEGFA antibodies"

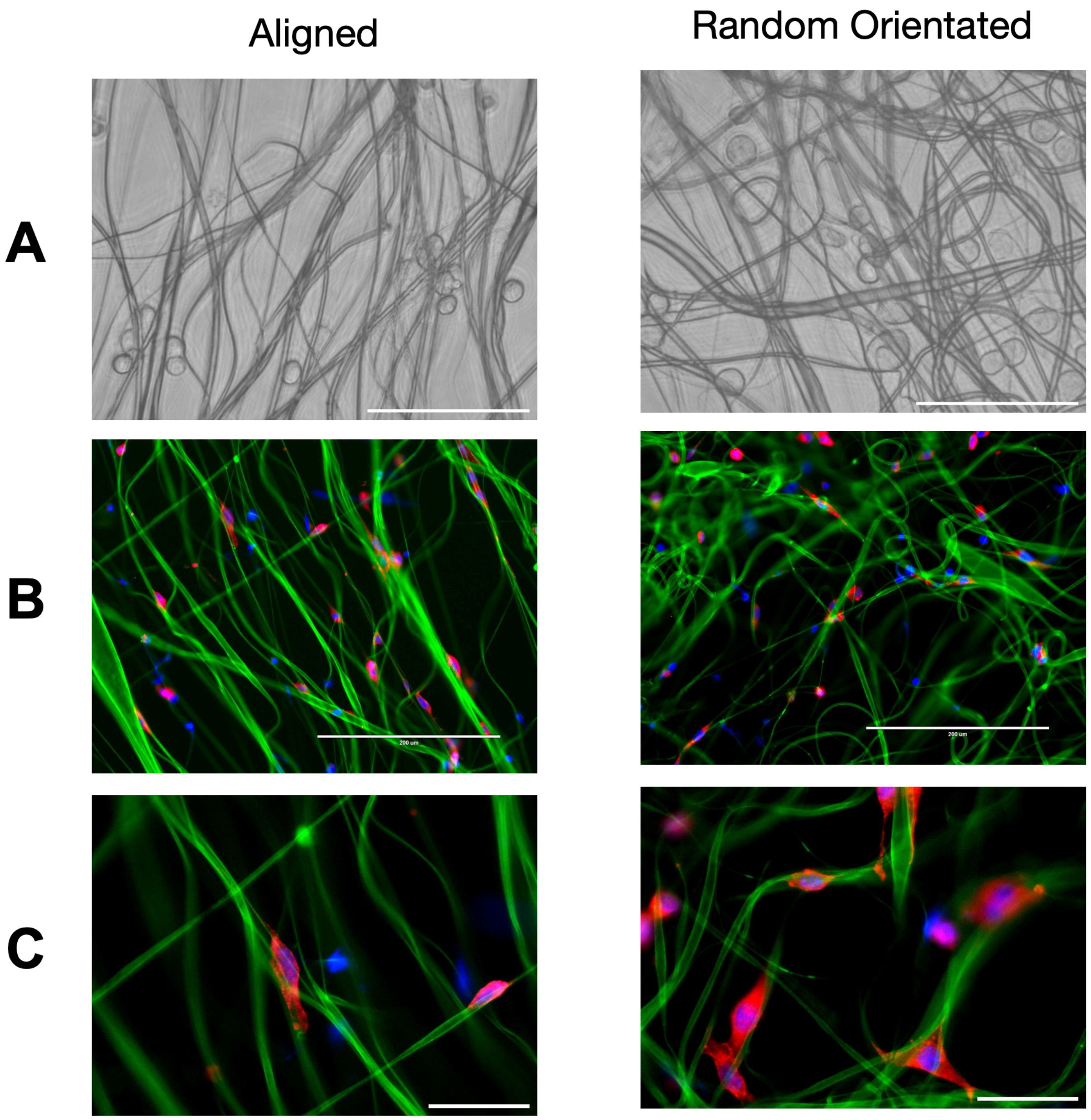


**Figure S2. (A)** Representative phase-contrast image of Fs120 cells cultured on fibronectin-coated fiber scaffolds for 24 hours (40X objectives). Scale bar = 100 µm. **(B and C)** Representative immunofluorescence images of Fs120 cells on fibronectin-coated fiber scaffolds after 24 hours, stained with phalloidin (red), fibronectin (green), and DAPI (blue). Images were captured using 20X **(B)** and 40X **(C)** objectives. Scale bars: 100 µm **(B)**, 50 µm **(C)**.
