## Supplemental Figure 3 for "Alternate VEGFA isoform expression in fibrosarcoma leads to plasticity in cellular migration and differences in sensitivity to inhibition with anti-VEGFA antibodies"

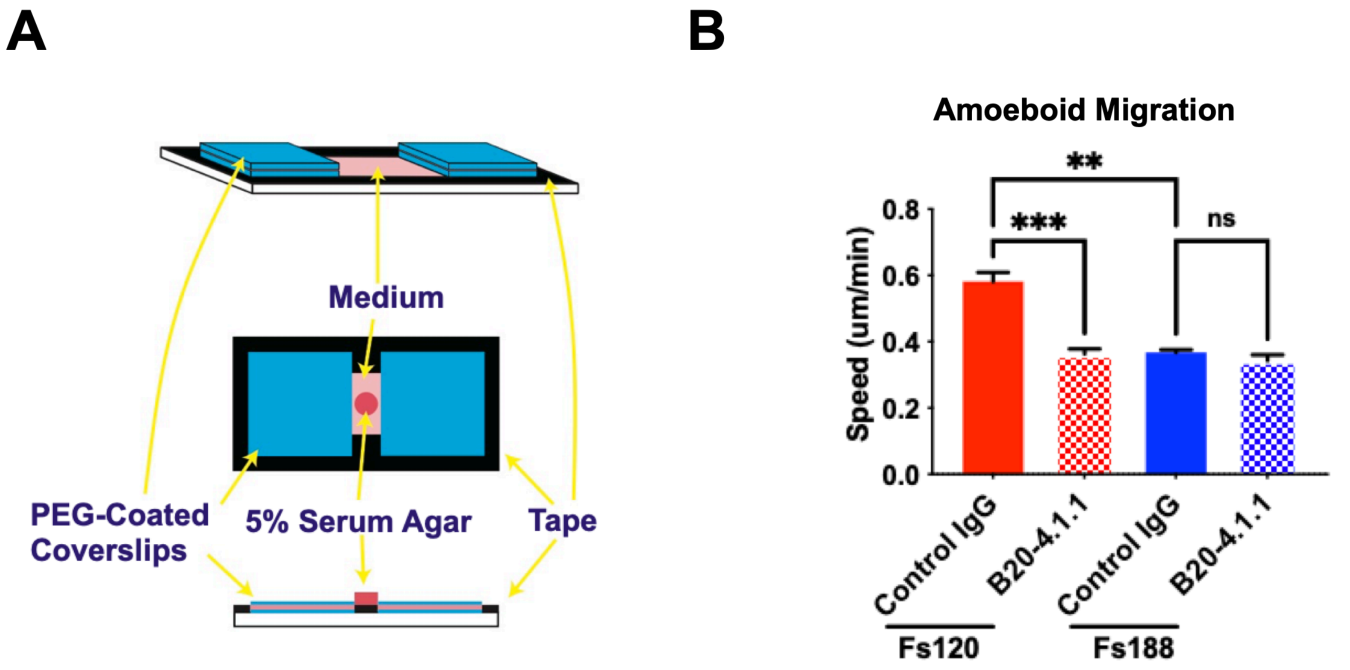


**Figure S3. (A)** The design of a non-adherent chamber was made by two cover glasses coated with PEG and spaced 40 μm apart with 3M tape on slides. The chemotactic gradient was facilitated by 5% (v/v) horse serum in agar. **(B)** Quantification of migration speed across fibrosarcoma cell lines in the non-adherent chamber from time-lapse imaging in the presence of 40 μg/mL B20-4.1.1 or its control IgG. Each column represents the mean of three independent replicates with error bars demoting ±SEM. Statistical significance (***p <0.001) and non-significance (ns) were determined using One-Way ANOVA with multiple comparisons.
