## Supplemental Figure 4 for "Alternate VEGFA isoform expression in fibrosarcoma leads to plasticity in cellular migration and differences in sensitivity to inhibition with anti-VEGFA antibodies"

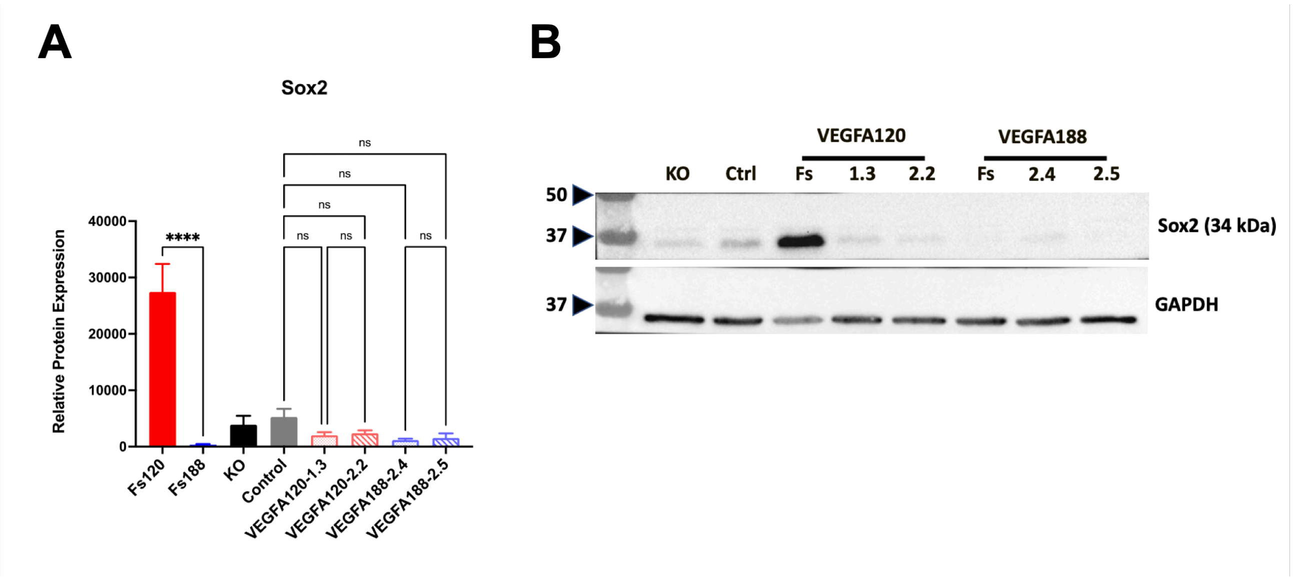


**Figure S4. (A)** Quantification of Sox2 protein relative expression in single VEGFA isoform expressing cells by Image J. Each column represents the mean of the three independent replicates with error bars demoting ±SEM. Statistical significance (****p <0.0001) and non-significance (ns) were determined using One-Way ANOVA with multiple comparisons by a Fisher's LSD post-test. **(B)** Representative Western blot images showing Sox2 expression in cells expressing individual VEGFA isoforms and VEGFA KO control cells are presented. GAPDH was used as a loading control. Western blot analysis was performed on three independent lysates.
