## Supplemental Figure 5 for "Alternate VEGFA isoform expression in fibrosarcoma leads to plasticity in cellular migration and differences in sensitivity to inhibition with anti-VEGFA antibodies"

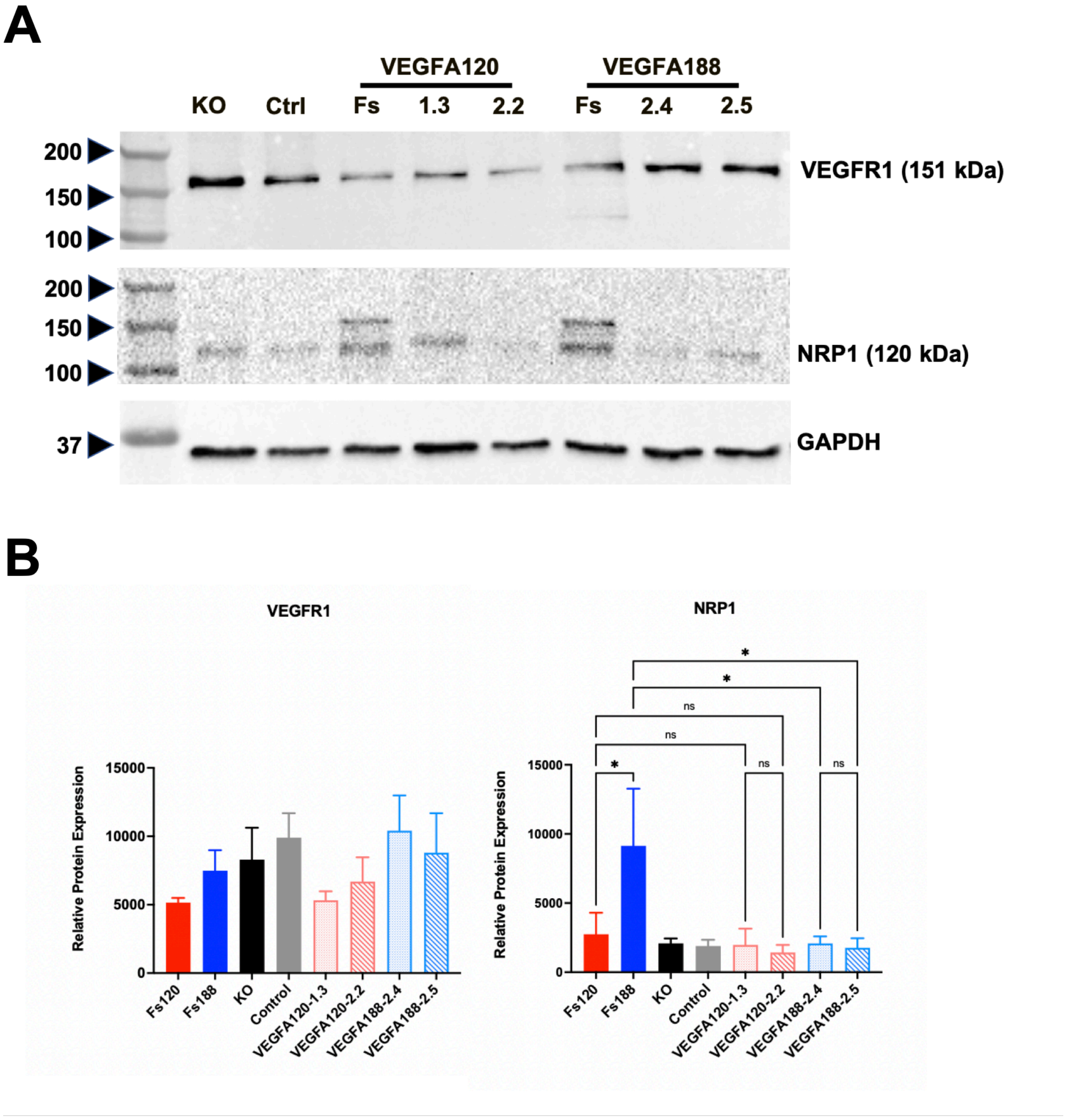


**Figure S5. (A)** Representative Western blot images showing Vegfr1 and Nrp1 expression in cells expressing individual VEGFA isoforms and VEGFA KO control cells are presented. GAPDH was used as a loading control. Western blot analysis was performed on three independent lysates. **(B)** Quantification of Vegfr1 and Nrp1 protein relative expression in single VEGFA isoform expressing cells by Image J. Each column represents the mean of the three independent replicates with error bars demoting ±SEM. Statistical significance (****p <0.0001) and non-significance (ns) were determined using One-Way ANOVA with multiple comparisons by a Fisher's LSD post-test.
